# The mouse Balbiani body maintains primordial follicle quiescence via RNA storage

**DOI:** 10.1101/2020.01.17.911040

**Authors:** Lei Lei, Kanako Ikami, Haley Abbott, Shiying Jin

**Author notes:** **Summary:** Integrity of the mouse Balbiani body maintains primordial follicle quiescence via mRNA-decapping enzyme 1A-mediated RNA storage. corresponding author: Lei Lei.

## Abstract

In mammalian females, the transition between quiescent primordial follicles and follicular development is critical for maintaining ovarian function and reproductive longevity. In primary oocytes of mouse quiescent primordial follicles, Golgi complexes are organized into a spherical structure, the Balbiani body. Here, we show that the structure of the B-body is maintained by microtubules and actin. The B-body stores mRNA-capping enzyme and 597 mRNAs associated with mRNA-decapping enzyme 1A. Proteins encoded by these mRNAs function in enzyme binding, cellular component organization and packing of telomere ends. Pharmacological disassembly of the B-body triggers translation of stored mRNAs and activates primordial follicles in culture and *in vivo* mouse model. Thus, primordial follicle quiescence is maintained by the B-body, and translationally inactive B-body-stored mRNAs may be regulated by 5’-capping.

## Main Text

In adult mammalian females, ovarian function is sustained by primordial follicles, each of which contains a primary oocyte and a single-layer of pregranulosa cells. Coordinated quiescence of primary oocytes and pregranulosa cells is essential for maintaining a pool of dormant primordial follicles, namely the ovarian reserve (*1*). A cohort of primordial follicles is periodically activated in the adult ovary to produce mature eggs (*2*). How primordial follicles remain quiescent and then a small cohort initiates development at a certain time remains a mystery. Mouse primary oocytes form by transferring cytoplasm and organelles from sister germ cells (*3*). In quiescent primary oocytes, the Golgi complexes together with centrosomes coalesce into a spherical structure, the Balbiani body (B-body), a highly conserved, oocyte-specific feature (*3, 4*) (Fig1.A, B). In zebrafish and *Xenopus*, the B-body, characterized by the presence of RNA granules and organelle aggregates (mitochondria and Golgi complex), is found in mature unfertilized eggs (*4, 5*). It plays an essential role in specifying germ cells in the early embryos via RNA storage and translational regulation. In postnatal mouse and human ovaries, the B-body is found only in quiescent primary oocytes, suggesting a potential connection between the B-body and primordial follicle quiescence (*6, 7*).

**Figure 1.**
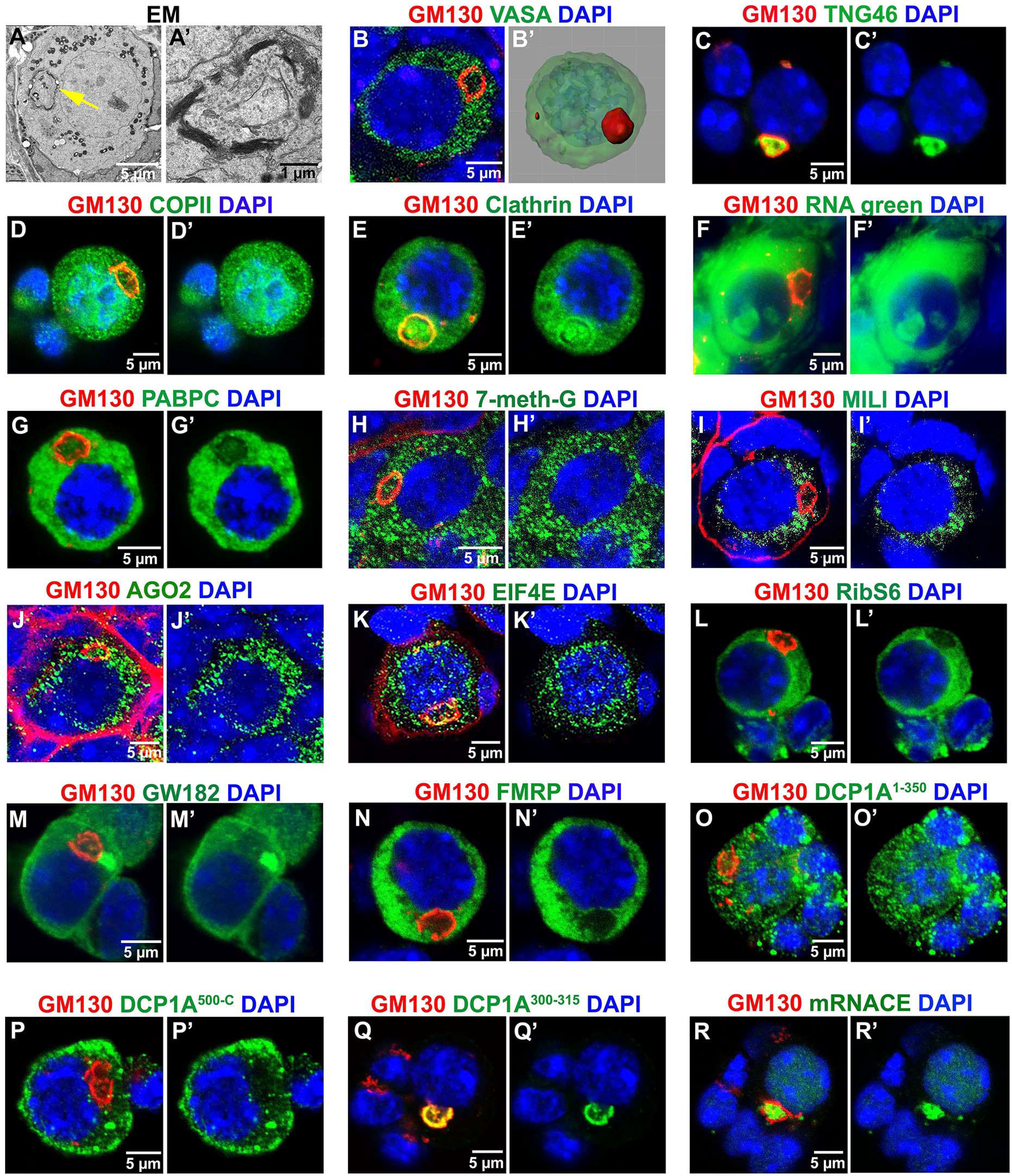
The B-body in quiescent primary oocytes characterized by EM images and antibody staining. (A) EM image showing a B-body (arrow) in a quiescent primary oocyte at lower magnification. (A’) A high magnification EM image showing highly stacked Golgi complexes organized in a circular structure. (B) A B-body stained by Golgi cis-face marker GM130 in a VASA positive quiescent primary oocyte. (B’) 3-D reconstructed image showing a B-body in a spherical structure. Co-immunostaining of the B-body by using GM130 and TNG46 (C), COPII (D), Clathrin (E), RNA green dye (F), PABPC (G), 7-methylguanosine (H), MILI (I), AGO2 (J), EIF4E (K), ribosomal protein S6 (L), GW182 (M), FMRP (N), DCP1A^1-350^(O), DCP1A^500-C^(P), DCP1A^300-315^ (Q), and mRNA capping enzyme (R).

To understand the nature and function of the mouse B-body, we determined how B-body Golgi complexes are organized together with other organelles in primary oocytes. We found that the trans-faces of the Golgi (marked by TNG46) are enclosed in the lumen of the B-body, and the cis-faces (marked by GM130) are exposed to the cytosol (fig1.C). Electron microscopy (EM) revealed many small vesicles residing inside and outside the B-body (Fig1A). We analyzed these vesicles by staining primary oocytes with a COPII antibody, which labels ER-to-Golgi transport vesicles, and with a Clathrin antibody, which marks secretary vesicles released from the Golgi trans-face (*8*). COPII-positive vesicles were found throughout the cytoplasm, but were relatively less abundant in the B-body (Fig1.D). By contrast, Clathrin-positive vesicles were enriched inside the B-body and also across the cis-face of the Golgi complexes (Fig.1E).

Enrichment of Trailer Hitch (TRAL), a protein linked to RNA metabolism, in the mouse B-body has led to the proposal that the B-body functions in RNA storage (*6*). To directly detect RNAs in the B-body, we stained primary oocytes with an RNA-binding dye. Strong signals in the cytoplasm and the nucleolus, but not in the rest of the nucleus, validated dye specificity. Compared with RNA levels in the cytoplasm, RNA content was lower in the B-body lumen (Fig.1F). Similarly, poly(A)-binding protein (PABPC), which interacts with mRNA poly(A) tails in the cytoplasm, was present at lower levels in the B-body lumen compared to the oocyte cytoplasm (Fig.1G) (*9, 10*). However, examining the distribution of mRNA 5’-caps, using the anti-7-methylguanosine antibody, revealed no obvious difference in 7-methylguanosine level or distribution either inside or outside the B-body (Fig.1H) (*11, 12*).

To elucidate whether or not the B-body contains small RNAs, we stained primary oocytes using antibodies to MILI (PIWI-like protein 2), which interacts with piRNAs, and Argonaute 2 (AGO2), a major component of the RNA-induced silencing complex (RISC) that represses translation by interacting with miRNAs and endo-siRNAs (*13*). MILI protein was excluded from the B-body, but otherwise distributed evenly in the cytoplasm (Fig. 1I). By contrast, AGO2 protein was found both inside and outside of the B-body, suggesting microRNAs but not piRNAs are a component of the B-body (Fig. 1J).

We asked whether proteins involved in translation were present within the B-body (*14*). Eukaryotic translation initiation factor 4E (EIF4E), required for translation initiation, was found both inside and outside of the B-body (Fig.1K). Ribosomal protein S6, required for elongation, was absent inside the B-body, suggesting that protein synthesis activity is low within the B-body (Fig.1L). The ribonucleoprotein (RNP) GW182, involved in miRNA-mediated translational silencing, was found in the cytoplasm and within the B-body, with a concentration of GW182 protein often found near the B-body (Fig.1M) (*15, 16*). Synaptic functional regulator FMR1 (FMRP), a polyribosome-associated RNA binding protein that mainly functions in repressing mRNA translation, was only found outside of the B-body in the cytoplasm (Fig.1N). mRNA-decapping enzyme 1A (DCP1A), a co-factor involved in removing the mRNA 5’ cap and thus facilitating RNA storage or degradation, showed distinct distribution patterns when using antibodies targeting different regions of the protein (*17*). The antibody targeting amino acids 1-350 detected small foci in the cytoplasm of primary oocytes and large foci in the surrounding pregranulosa cells (Fig.1O). The antibody targeting amino acid 500 to the C-terminus of the protein detected several small granules in both primary oocytes and pregranulosa cells (Fig. 1P). The antibody targeting amino acids 300-315 specifically detected protein located on the inner side of the Golgi cis-face of the primary oocyte (Fig.1Q). Like DCPA1^300-315^, the mRNA-capping enzyme (mRNACE) was highly enriched inside of the B-body (Fig.1R). These observations suggest that the B-body may be involved in RNA storage and translational regulation, possibly via 5’-cap modifications of mRNAs.

The B-body is only found in quiescent primary oocytes (Fig.1A). EM images revealed that in the activated oocytes, the spherical structure of the B-body was disassociated (Fig.2A). To investigate how the integrity of the B-body is maintained, we examined the distribution of cytoskeleton proteins in quiescent primary oocytes (*18, 19*). We found that α-tubulins were highly enriched around the B-body and co-localize with the Golgi cis-face, suggesting a close interaction between microtubules and Golgi cis-face (Fig.2B). By contrast, stable microtubules, represented by acetylated tubulins, did not directly co-localize with the Golgi complexes, but in a network around the B-body (Fig.2C). Despite enriched tubulins around the B-body, microtubules were not aligned in a polarized manner in the primary oocyte. Microtubule minus-end binding protein CAMSAP3 and plus-end tracking protein EB1 were evenly distributed in the primary oocyte and could be detected both inside and outside of the B-body (Fig.2D, E)(*20, 21*). F-actin, detected by phalloidin, was observed to be heavily distributed along the primary oocyte cell membrane and in several clusters around the B-body (Fig.2F). We carried out a comparative proteomic analysis of quiescent primordial follicles and developing follicles. 35 actin-related proteins were identified, 24 of these proteins were more enriched in primordial follicles than in developing follicles. 31 tubulin-related proteins were identified and only four proteins were more abundant in primordial follicles (Fig.2G, Supp Table 1). These results suggest that microtubules and actin are responsible for maintaining B-body structure in quiescent primary oocytes.

**Figure 2.**
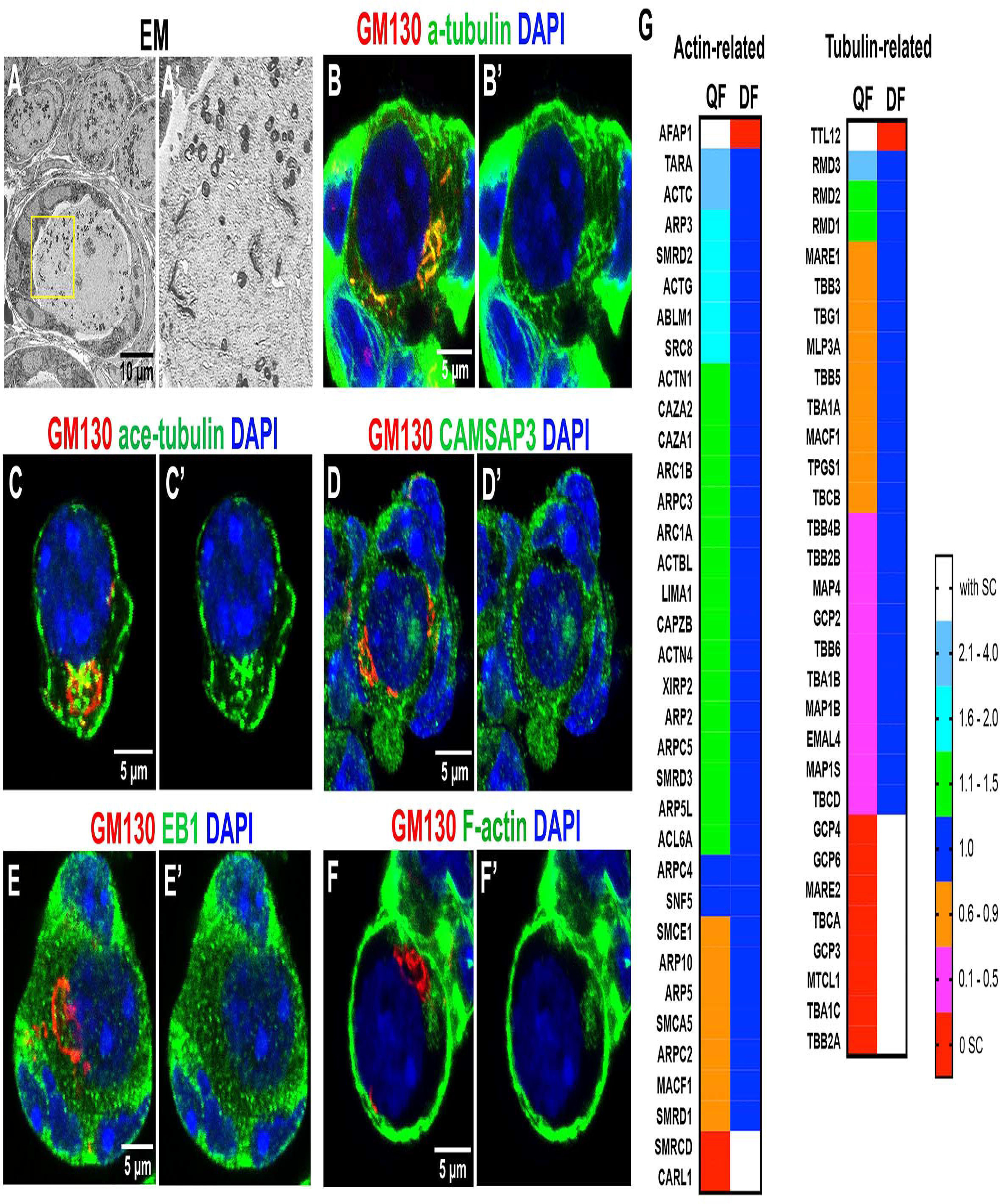
Actin and microtubules are involved in B-body integrity. (A) EM images showing a disassociated B-body in an activated follicle. (B-F) distribution of α-tubulin (B), acetylated-tubulin (C), CAMSAP3 (D), EB1 (E), and F-actin (F) in quiescent primary oocytes. (G) Fold change of normalized spectral abundance factor of actin-related proteins and tubulin-related proteins in quiescent primordial follicles (QF) and developing follicles (DF), SC (spectral count).

To test directly the role of microtubules and actin in maintaining B-body integrity, and to elucidate the causative connection between B-body integrity and primordial follicle activation, we first used an *in vitro* ovarian culture model to manipulate microtubule or actin dynamic within cells. postnatal day 4 (P4) ovaries, in which over 90% follicles are quiescent primordial follicles were dissected, incubated with polymerization inhibitors of actin (cytochalasin D at 10 μM) or tubulin (nocodazole at 10 μM), then cultured for 6 days to allow activated follicles to grow (Fig.3A). The percentages of primary oocytes containing a B-body in control, cytochalasin D-treated, or nocodazole-treated ovaries were 85.9%, 44.6%, and 29.5%, respectively, indicating both inhibitors significantly disrupted B-body integrity (Fig.3B, C). In cytochalasin D-treated primary oocytes, the Golgi complexes remained linear and clustered, but had lost its spherical structure. In nocodazole-treated primary oocytes, Golgi complexes fragmented into small pieces and dispersed throughout the primary oocyte (Fig.3B). This result suggests that microtubules and actin may function differently in maintaining B-body integrity. To understand the effect of B-body disassociation, we analyzed ovarian morphology after a 6-day culture and a significantly increased number of developing follicles, recognized by the larger volume of their VASA+ oocytes, were observed in drug-treated ovaries (Fig.3D). On average, 65.2±17.9 developing follicles per ovary were found in control, versus 154.8±11.7 in cytochalasin D-treated ovaries and 124.4±20.5 in nocodazole-treated ovaries (Fig.3E). Similar effects of follicle activation were also observed when we injected cytochalasin D or nocodazole to P4 mice, about two times more developing follicles were found in ovaries of injected mice than control mice (Fig.3A, F). These results demonstrate that B-body disassociation activates primordial follicle.

**Figure 3.**
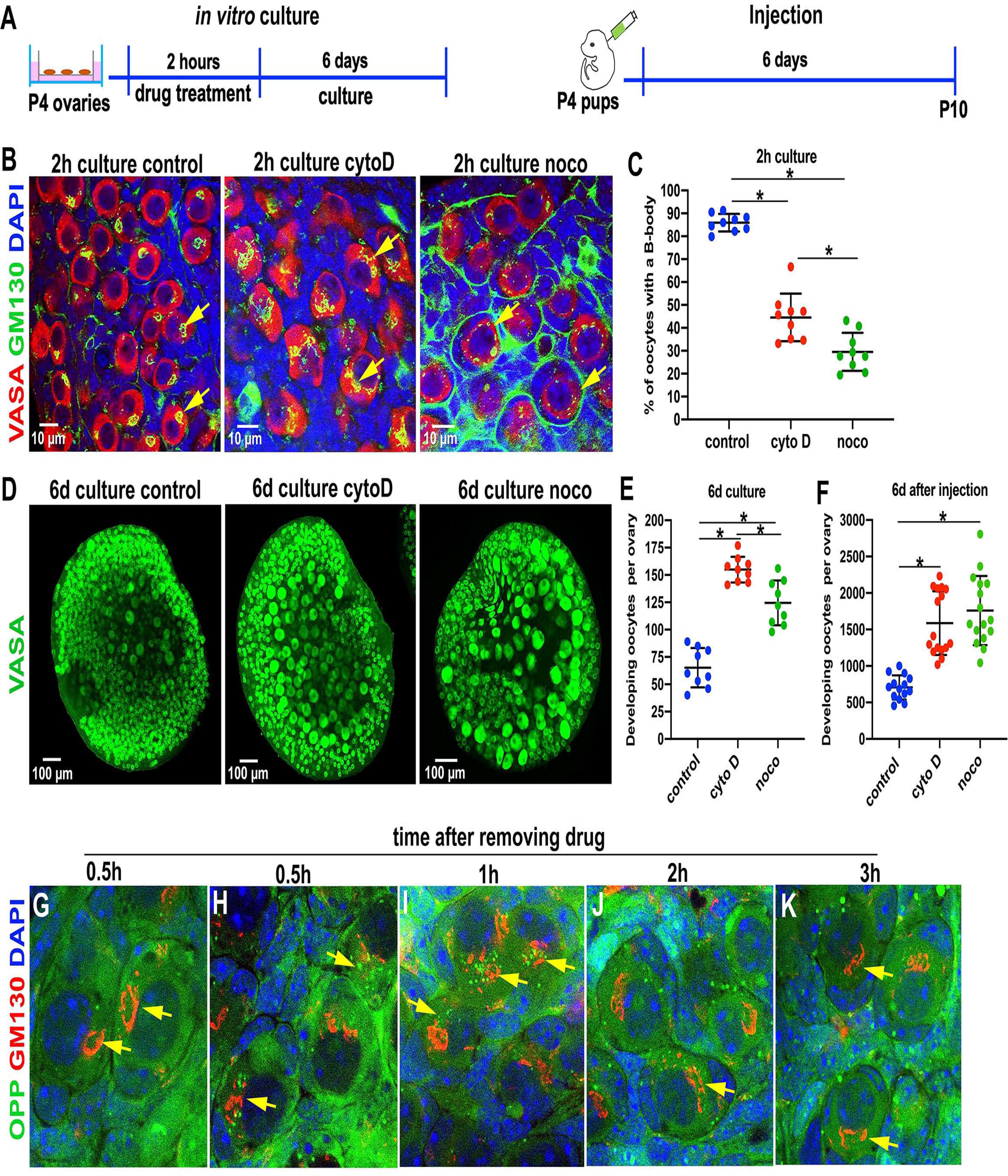
Disassociating the B-body activates primordial follicles. (A) scheme of ovarian culture and mouse injection for primordial follicle activation. (B) Immunostaining of GM130 showing the Golgi complex morphology after drug treatment. (C) Percentage of primary oocytes containing a B-body in the ovaries after a 2-hour drug treatment. (D) Oocyte morphology of drug-treated ovaries after a 6-day culture. (E) Number of developing follicles in each drug-treated ovary after a 6-day culture. (F) Number of developing follicles in each P10 ovary after a drug injection. Results were analyzed statistically by using one-way ANOVA. P<0.05 was considered to be statistically significant *. (G-K) Nascent protein synthesis revealed by fluorescent positive Opropargyl-puromycin (OPP) foci in the ovaries treated with Cytochalasin D for 2 hours.

To reveal whether protein synthesize takes place after B-body disassociation, O-propargyl-puromycin (OPP), a marker for newly synthesized protein, was added to cultured ovaries at various times following incubation of cytochalasin D. After 0.5 h, OPP-positive protein foci were observed near disassociated B-bodies but were absent around the intact spherical B-bodies at all time points (Fig.3F, G, Supp Fig.1). Protein foci appeared largest around disassociated B-bodies 1h after removal of cytochalasin D (Fig.3H). At 2h, foci were more dispersed in the cytoplasm, and by 3h, almost no foci were observed in the primary oocytes with disassociated B-bodies (Fig.3I, J). Thus, local protein synthesis, which takes place within four hours of B-body disassociation may promote the activation of primordial follicles.

To identify RNAs present in the B-body, we conducted RNA-immunoprecipitation (RIP) by using the DCP1A^300-315^antibody (Supp Fig.2). 597 mRNAs were significantly enriched in DCP1A pull-down RNA samples and thus were abundant in the B-body. The majority (451 out of 597, 75.5%) of these mRNAs encoded proteins were absent in quiescent primordial follicles, confirming that most B-body-associated RNAs are not translated in primordial follicles. The most represented molecular function (16 in total), biological process (56 in total) and REACTOM (61 in total) of these mRNAs are enzyme binding, cellular component organization and packaging of telomere ends (Fig.4C-E). Collectively, these data revealed the initial cellular and molecular processes taking place upon B-body disassociation that may promote primordial follicle activation.

**Figure 4.**
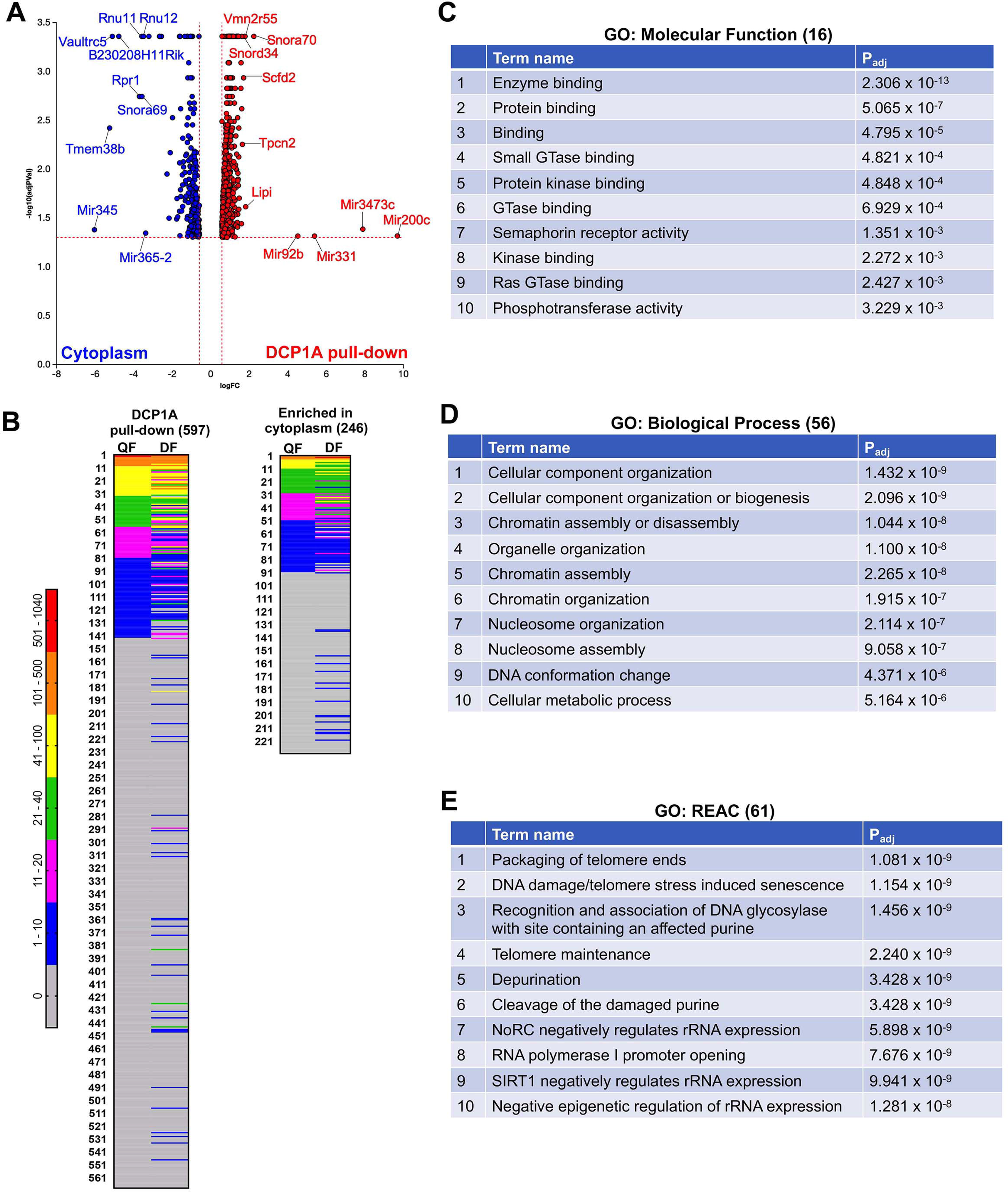
Profiles of the B-body stored mRNAs enriched by RNA immunoprecipitation with DCP1A. (A) Volcano plot showing the enriched mRNAs in the B-body or cytoplasm. (B) The protein expression (normalized spectral count) of the DCP1A-pull down mRNAs and cytoplasm-enriched mRNAs in quiescent primordial follicles (QF) and developing follicles (DF). Gene ontology analysis results of the top 10 candidates of the molecular function (C), biological process (D), REACTOM (E) of DCP1A-pull down mRNA.

Here, we show that the mouse B-body maintains primordial follicle quiescence via RNA storage by using DCP1A. While organelle aggregates also facilitate translational regulation in oocytes in zebrafish and frogs, our study suggests that a distinct 5’-cap modification-related mechanism is used in mice for mRNA storage and translational regulation. The close association of mRNA-capping enzyme with DCP1A in the disassociated B-body suggests that translational regulation of the DCP1A-associate mRNA is facilitated through the addition of mRNA 5’-caps, a process identified in hepatitis B virus and in erythroid cells of a beta-thalassemic mouse model (Supp Fig.5) (*22*). We show that B-body integrity, maintained by actin and microtubules, is critical for primordial follicle quiescence, with activation of primordial follicles requiring disruption of B-body structure. The human B-body has been identified as a cluster of organelles, including centrosomes, Golgi complexes, and mitochondria next to the nucleus(*7*). Whether a spherical Golgi complex forms part of the human B-body is not known, although the clustered organelles that distinguish the B-body is similar to the mouse B-body (*23*). Primordial follicles quiescence may be maintained by a similar mechanism in mice and humans, given that PTEN inhibitors have been shown to activate mouse and human quiescent primordial follicles (*24, 25*). The novel cellular and molecular processes of primordial follicle activation uncovered in the present study shed lights on mechanisms of fertility loss due to follicle activation upon chemotherapy. Our study also reveals a potential medical application of pharmacologically manipulating the B-body integrity to preserve or activate primordial follicles for the purposes of fertility preservation, ovarian longevity and infertility treatment.

## Acknowledgements

Research reported in this paper was supported by the Sergey Brin Family Foundation.

## References

1. M. E. Pepling, Follicular assembly: mechanisms of action. Reproduction (Cambridge, England) 143, 139–149 (2012).

2. S. Lintern-Moore, G. P. Moore, The initiation of follicle and oocyte growth in the mouse ovary. Biology of reproduction 20, 773–778 (1979).

3. L. Lei, A. C. Spradling, Mouse oocytes differentiate through organelle enrichment from sister cyst germ cells. Science 352, 95–99 (2016).

4. M. Kloc, S. Bilinski, L. D. Etkin, The Balbiani body and germ cell determinants: 150 years later. Current topics in developmental biology 59, 1–36 (2004).

5. E. Boke et al., Amyloid-like Self-Assembly of a Cellular Compartment. Cell 166, 637–650 (2016).

6. M. E. Pepling, J. E. Wilhelm, A. L. O’Hara, G. W. Gephardt, A. C. Spradling, Mouse oocytes within germ cell cysts and primordial follicles contain a Balbiani body. Proceedings of the National Academy of Sciences of the United States of America 104, 187–192 (2007).

7. A. T. Hertig, The primary human oocyte: some observations on the fine structure of Balbiani's vitelline body and the origin of the annulate lamellae. Am J Anat 122, 107–137 (1968).

8. J. Klumperman, Architecture of the mammalian Golgi. Cold Spring Harbor perspectives in biology 3, (2011).

9. S. A. Adam, T. Nakagawa, M. S. Swanson, T. K. Woodruff, G. Dreyfuss, mRNA polyadenylate-binding protein: gene isolation and sequencing and identification of a ribonucleoprotein consensus sequence. Molecular and cellular biology 6, 2932–2943 (1986).

10. C. G. Burd, E. L. Matunis, G. Dreyfuss, The multiple RNA-binding domains of the mRNA poly(A)-binding protein have different RNA-binding activities. Molecular and cellular biology 11, 3419–3424 (1991).

11. U. Schibler, R. P. Perry, Characterization of the 5' termini of hn RNA in mouse L cells: implications for processing and cap formation. Cell 9, 121–130 (1976).

12. D. R. Schoenberg, L. E. Maquat, Re-capping the message. Trends in biochemical sciences 34, 435–442 (2009).

13. D. J. Gibbings, C. Ciaudo, M. Erhardt, O. Voinnet, Multivesicular bodies associate with components of miRNA effector complexes and modulate miRNA activity. Nature cell biology 11, 1143–1149 (2009).

14. N. K. Gray, M. Wickens, Control of translation initiation in animals. Annual review of cell and developmental biology 14, 399–458 (1998).

15. T. Eystathioy et al., A phosphorylated cytoplasmic autoantigen, GW182, associates with a unique population of human mRNAs within novel cytoplasmic speckles. Molecular biology of the cell 13, 1338–1351 (2002).

16. T. Eystathioy et al., A panel of monoclonal antibodies to cytoplasmic GW bodies and the mRNA binding protein GW182. Hybridoma and hybridomics 22, 79–86 (2003).

17. J. Coller, R. Parker, Eukaryotic mRNA decapping. Annual review of biochemistry 73, 861–890 (2004).

18. J. Thyberg, S. Moskalewski, Role of microtubules in the organization of the Golgi complex. Experimental cell research 246, 263–279 (1999).

19. K. Heimann, J. M. Percival, R. Weinberger, P. Gunning, J. L. Stow, Specific isoforms of actin-binding proteins on distinct populations of Golgi-derived vesicles. The Journal of biological chemistry 274, 10743–10750 (1999).

20. W. Meng, Y. Mushika, T. Ichii, M. Takeichi, Anchorage of microtubule minus ends to adherens junctions regulates epithelial cell-cell contacts. Cell 135, 948–959 (2008).

21. L. Berrueta et al., The adenomatous polyposis coli-binding protein EB1 is associated with cytoplasmic and spindle microtubules. Proc Natl Acad Sci U S A 95, 10596–10601 (1998).

22. J. B. Trotman, D. R. Schoenberg, A recap of RNA recapping. Wiley interdisciplinary reviews. RNA 10, e1504 (2019).

23. A. T. Hertig, E. C. Adams, Studies on the human oocyte and its follicle. I. Ultrastructural and histochemical observations on the primordial follicle stage. The Journal of cell biology 34, 647–675 (1967).

24. M. McLaughlin, H. L. Kinnell, R. A. Anderson, E. E. Telfer, Inhibition of phosphatase and tensin homologue (PTEN) in human ovary in vitro results in increased activation of primordial follicles but compromises development of growing follicles. Molecular human reproduction 20, 736–744 (2014).

25. J. Li et al., Activation of dormant ovarian follicles to generate mature eggs. Proceedings of the National Academy of Sciences of the United States of America 107, 10280–10284 (2010).

